# Uncovering genetic architecture of the heart via genetic association studies of unsupervised deep learning derived endophenotypes

**DOI:** 10.1101/2025.09.17.676827

**Authors:** Lei You, Xingzhong Zhao, Ziqian Xie, Khush A Patel, Cheng Chen, Danai Kitkungvan, Kamil Khan Mohammed, Navneet Narula, Eloisa Arbustini, Craig K. Cassidy, Jagat Narula, Degui Zhi

## Abstract

Recent genome-wide association studies (GWAS) have effectively linked genetic variants to quantitative traits derived from time-series cardiac magnetic resonance imaging, revealing insights into cardiac morphology and function. Deep learning approach generally requires extensive supervised training on manually annotated data. In this study, we developed a novel framework using a 3D U-architecture autoencoder (cineMAE) to learn deep image phenotypes from cardiac magnetic resonance (CMR) imaging for genetic discovery, focusing on long-axis two-chamber and four-chamber views. We trained a masked autoencoder to develop **U**nsupervised **D**erived **I**mage **P**henotypes for heart (Heart-UDIPs). These representations were found to be informative to indicate various heart-specific phenotypes (e.g., left ventricular hypertrophy) and diseases (e.g., hypertrophic cardiomyopathy). GWAS on Heart UDIP identified 323 lead SNP and 628 SNP-prioritized genes, which exceeded previous methods. The genes identified by method described herein, exhibited significant associations with cardiac function and showed substantial enrichment in pathways related to cardiac disorders. These results underscore the utility of our Heart-UDIP approach in enhancing the discovery potential for genetic associations, without the need for clinically defined phenotypes or manual annotations.

## Introduction

Cardiac magnetic resonance imaging (CMR) has become one of the most commonly utilized imaging modality for the heart. A non-contrast enhanced CMR cine imaging allows detailed assessment of cardiac anatomy and function at high spatiotemporal resolution.

With the availability of large cohorts that include both CMR data and genetic information[1-3], investigators can identify genetic variations influencing individual differences in cardiac structure and function through the analysis of parallel genome-wide association studies (GWAS) from the same cohorts. Such genetic insights may enhance our understanding of various cardiac pathologies, such as ventricular dilation and dysfunction observed in dilated cardiomyopathy (DCM)[4], myocardial hypertrophy characteristic of hypertrophic cardiomyopathy (HCM)[5] or remodeling following myocardial infarction in ischemic heart disease (IHD)[1, 6].

One of the main challenges of GWAS in heart-imaging is the derivation of comprehensive, heritable, and interpretable representations of the heart from complex multi-dimensional CMR data. Most existing GWAS of cardiac phenotypes rely on classical measures such as left ventricular end-diastolic volume (LVEDV), left ventricular end-systolic volume (LVESV), ejection fraction (EF), and myocardial mass, often computed with semi-automated or manual approaches[3]. While newer deep learning pipelines have streamlined the derivation of such conventional metrics, they typically focus on automating existing measurements rather than discovering entirely new phenotypic dimensions[7].

However, traditional CMR phenotypes face inherent limitations. Initially, deriving standard metrics is time-consuming and potentially inconsistent, as they often rely on manual or semi-automated segmentations that introduce user and software bias and are limited by pre-existing human knowledge. Moreover, restricting analysis to a few global measures (e.g., EF or LV volumes) can fail to capture subtle but genetically informative structural and functional variations within the myocardium. Single measures, such as EF, aggregate information from distinct myocardial regions into one summary statistic, potentially obscuring localized pathology or genetic influences that are spread across multiple cardiac segments. Some multivariate statistical approaches have attempted to jointly model multiple phenotypes, thereby improving sensitivity to genetic signals. However, these still require a priori determination of which phenotypes to include, risking the exclusion of equally relevant but unmeasured aspects of cardiac structure and function (van der Meer et al., 2020[8], Guo B et al.[9]).

Our previous work addressing brain MRI [10] has enabled direct extraction of imaging phenotypes from imaging data without predefined labels. We developed an **U**nsupervised **D**erived **I**mage **P**henotypes (UDIP) tool that uses an autoencoder with a bottleneck layer and is trained by a voxel-wise reconstruction loss. The bottleneck layers are a compact representation of the input because they contain essential information for accurately representing the input. We use the activation of the bottleneck layer as an endophenotype of the input image. The advantage is that UDIP requires minimal preprocessing and no human labeling. We demonstrated the initial success of the UDIP approach for brain MRI images. Our method yields phenotypes with higher heritability as defined below (heritability of Heart-UDIPs), leading to the discovery of novel genetic loci from biobank-scale datasets. In this work, we aim to develop the UDIP framework for heart imaging data.

While the conceptual idea of UDIP seems generalizable [11], several challenges remain for CMR due to significant differences in brain and heart images. First, unlike the brain, which is the only organ encapsulated in the skull, the heart shares the thorax with other organs, and thus, a careful segmentation is needed. Second, unlike static brain MRI, CMR data capture a beating heart, requiring modeling across cardiac cycles. Moreover, the orientation and geometry of the heart vary significantly among individuals, and image registration becomes important. While many atlases exist for brain MRI, no universal cardiac coordinate system currently exists. These obstacles revealed essential components that were missing in previous UDIP approaches.

Therefore, we developed a general deep learning model to extract Heart Unsupervised Derived Image Phenotypes (Heart-UDIP), applicable to both 2Ch and 4Ch cardiac MR images. This interpretable Heart-UDIP captured useful morphological and functional information about the heart. We demonstrated that these Heart-UDIPs serve as powerful biomarkers, facilitating the discovery of distinct signatures for heart diseases such as hypertrophic and dilated cardiomyopathy in both imaging and genetic analysis. We found many novel heart-related genes through GWAS of our Heart-UDIPs. To support this analysis, we developed an essential preprocessing pipeline that incorporates two cine MRI atlas image registration and segmentation across different views.

## Results

### Overview

This study aims to advance our understanding of cardiomyopathies by moving beyond the traditional, simple cardiac measurements through the use of unsupervised learning, which enables the interrogation of CMR images with unprecedented detail and granularity. A key innovation is the development of a unified deep learning model, capable of processing both the 2Ch and 4Ch views, enabling view-specific representation of the cardiac anatomy and function. Our genetic discovery underscores the power of this approach; an analysis of the 2Ch view phenotypes identified 181 lead SNPs, while the 4Ch view analysis yielded 180 lead SNPs. Combining the results allowing us to prioritize 628 genes and uncover novel biological pathways related to myocardial diseases. Our workflow involved developing these deep learning models, extracting novel phenotypes, and performing subsequent genetic and biological pathway analyses.

We analyzed CINE CMR images from an initial cohort of 77,050 UK Biobank participants. For deep learning models, we first selected 10,000 individuals without a history of heart failure to serve as the training dataset. The remaining 67,050 participants were designated as the test set for the network training and GWAS. The test set underwent an initial quality control (QC) step where we excluded individuals whose images failed our automated segmentation and registration pipeline, resulting in 66,874 participants for phenotype extraction. For the subsequent GWAS study, subjects with missing covariates were dropped (**Methods**), yielding a final analysis cohort of 58,405 individuals. Both 2Ch and 4Ch view images were used throughout the experiments. Further details on the cohort selection and QC criteria are provided in the Methods section. More details can be found at **Figure S1**.

We designed a new efficient architecture to extract the UDIPs from those CMR images. The architecture features a 3D convolutional autoencoder with a central 256-dimensional bottleneck layer. To improve reconstruction fidelity and reduce image blur, we incorporate a U-Net-like skip connection that links the final encoder layer to the initial decoder layer (**Figure 1b, Methods**). Unlike brain UDIP, we trained the model with masked input for better representation learning. For each subject, the 256-dimensional feature vector from the bottleneck layer served as the set of derived phenotypes (or Heart-UDIP).

**Figure 1.**
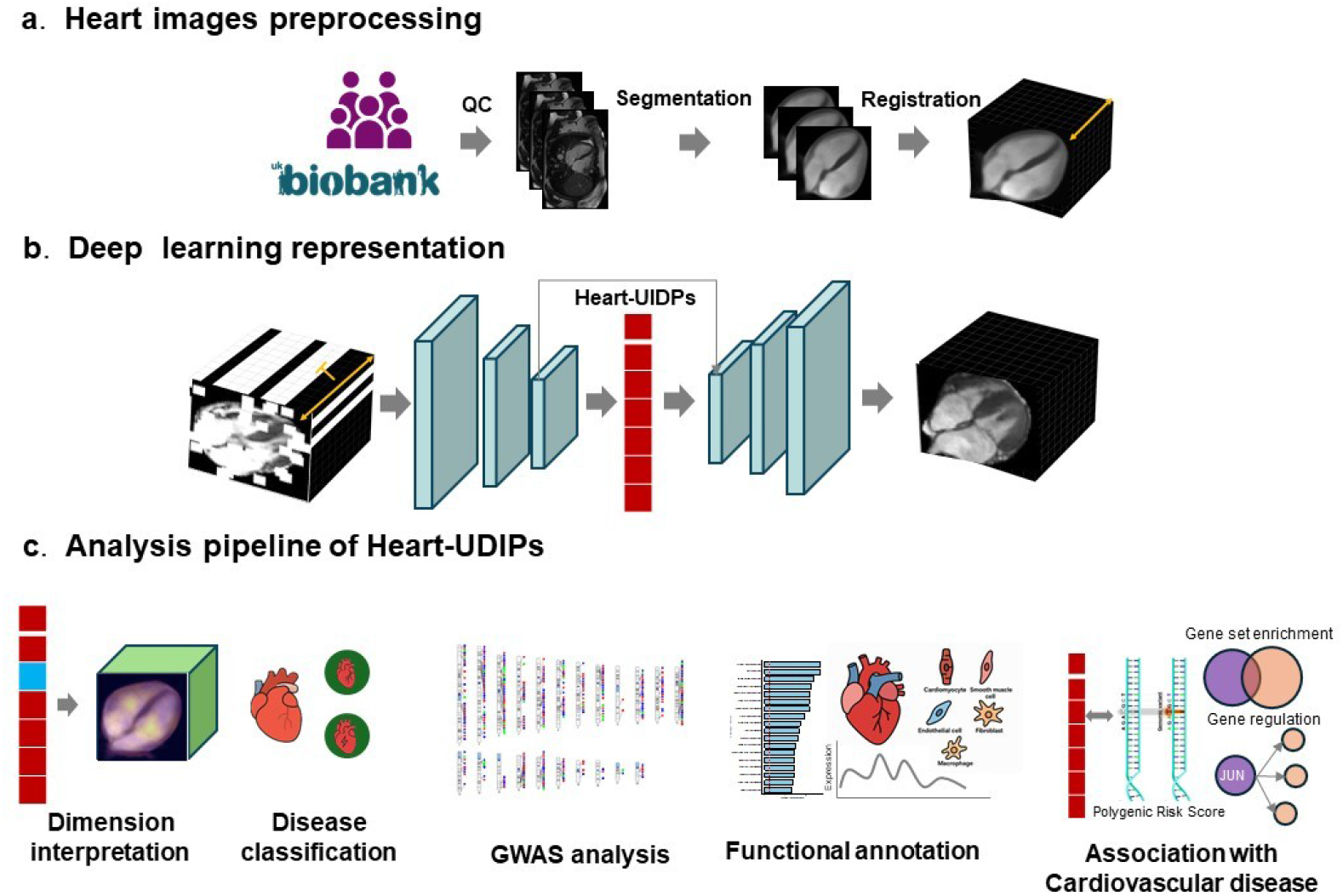
The workflow of our study. **(a)** Pipeline of the preprocessing for the CMR images. **(b)** We proposed a UNet-like mask autoencoder (**cineMAE**) which is utilized to extract the image phenotypes from the Cine MR images. We named those phenotypes Heart-UDIPs. **(c)** Analysis pipeline for the Heart-UDIPs.

To evaluate the interpretability and biological relevance of Heart-UDIPs, we conducted analyses that (i) map latent dimensions to specific cardiac anatomy, (ii) evaluated their performance in disease classification, (iii) identified the genetic architecture of heart through GWAS of Heart-UDIPs, and (iv) revealed the connections of these phenotypes with cardiovascular disease risk through gene-set enrichment, regulatory network analysis, and polygenic risk scoring. Our analyses established a link from imaging signal to genetic architecture and clinical outcomes (**Figure 1c**).

### Image data utilized in this study

Our image dataset comprises imaging data from 77,050 participants in the UK Biobank. The CMR images were available as 2Ch, 3Ch, and 4Ch long-axis views. For each view, the images are taken at a fixed position of the heart by 50 frames. In our experiment, both 2Ch view and 4Ch view CMR images are selected for image feature extraction and GWAS analysis as they provide orthogonal views of cardiac anatomy. The resolution of those CMR images is 208*168. We randomly selected 10,000 patients without any heart failure-related ICD-10 codes to form the training set. The remaining cases were designated for testing. After excluding cases with missing data or failed segmentation, 66,874 cases remained in the image feature related test set. There is no overlap between the training and test sets.

To isolate the heart and standardize anatomical alignment, we developed a two-stage preprocessing pipeline involving segmentation and registration. First, to exclude irrelevant information from surrounding organs, we trained a nnUNet[12] model to segment the heart from both 2Ch and 4Ch view images. This model was trained on a small, annotated dataset of 100 randomly selected participants for each view. During inference, any participant whose heart could not be successfully segmented by the nnUNet was excluded from analysis.

Second, after confirming the importance of spatial alignment for genetic discovery, we addressed the lack of a public cardiac MRI atlas for the UK Biobank. We created a novel heart atlas by selecting 150 of the successfully segmented images from 2Ch and 4Ch views and registered them to a common space. The resulting atlas has a resolution of 80x80x50 voxels. All images in our study were subsequently registered to this atlas before being used for model training, ensuring anatomical consistency across the entire cohort (**Methods**).

### cineMAE: cine MR Image Masked Autoencoder

To enable quantitative phenotype generation for genetic discovery, we introduce cineMAE, a masked autoencoder specifically designed to learn high-fidelity representations from cardiac cine MRIs. The model employs a 3D convolutional encoder–decoder architecture with a central 256-dimensional bottleneck layer, where each encoder block integrates 3D convolutions, ReLU activations, and instance normalization to effectively capture spatial-temporal structure (**Methods**).The model was trained using a loss function that combines a mean square error (MSE) reconstruction loss on masked images with a structural similarity index measure (SSIM) loss to ensure high-fidelity reconstruction of anatomical details. The resulting 256-dimensional bottleneck vectors constitute the Heart-UDIPs. For genetic discovery, we conducted per-UDIP GWAS using a Bonferroni-corrected genome-wide threshold of 5e-8/256.

A key architectural innovation is the addition of a skip-connection between the deepest layers of the autoencoder. This simple addition significantly enhances reconstruction quality, producing sharp images free of blurring artifacts, even with an 80% input masking ratio. This representation proved critical for downstream success. In our early experiments using array genotype data, the model without skip connections failed to identify any significant SNPs,whereas the model with skip-connection produced features showing genome-wide significant associations (**Supplementary Table 1**).

A second key innovation of our work is the use of a masking strategy during training, which forces the model to learn more discriminative and holistic representations from the highly redundant data in cine CMR. Long-axis cine CMR sequences contain substantial spatial redundancy, as adjacent 2D frames are often nearly identical. Applying standard 3D convolutions to this data results in inefficient parameter utilization. To address this, we implemented a masking strategy that crops a significant portion of the input, compelling the model to focus on learning essential spatio-temporal features rather than redundant spatial information. Specifically, we applied a tube mask with a size of 8x8x50. The Heart-UDIP generated using this masking strategy demonstrated superior disease classification performance (**Supplementary Table 2**). For dilated cardiomyopathy, the area under the curve (AUC) increased from 0.75 with standard IDP to 0.84 using IDP from cineMAE. The greater improvements in 4Ch-view indicate that the masking strategy brings more benefits to the models trained on the views with richer structured information.

Applying masking during representation learning increased the yield of genome-wide significant associations. Relative to unmasked models, masked Heart-UDIP identified 116 additional lead SNP for 2Ch-derived phenotypes and 73 for 4Ch-derived phenotypes (**Supplementary Table 3**). The masked representations also exhibited lower inter-feature correlation, indicating more disentangled dimensions. Consequently, the resulting phenotype set is more discriminative and information-rich, increasing power for genetic discovery.

### Ablation results

Since our main aim was to find cardiac-related significant SNPs from our Heart-UDIP, we used the number of SNPs found as the evaluation criteria to show how those preprocessing steps and masking strategy influenced the results. All the ablation experiments were conducted using array genotype for computational efficiency. **Supplementary Table 3** shows that both image segmentation and image registration boost genetic discovery. No significant SNPs were found by the traditional 3DCNN models. In contrast, cineMAE identified two SNPs even without image segmentation and registration. Adding image segmentation to the cineMAE increased this to five SNPs, while incorporating both image segmentation and registration yielded seven SNPs. The largest boost in the number of discovered SNP came from the masking strategy, increasing the number to 80. These results show that our preprocessing pipeline helps the model to learn more informative Heart-UDIPs that can lead to more SNP discovery.

### Image reconstruction quality

To evaluate the success of training, we randomly picked 5000 patients from the test dataset with both 2Ch and 4Ch images and calculate the mean Structural Similarity Index (SSIM) and mean peak-signal-to-noise ratio (PSNR), between the original images and their corresponding reconstructions from the model. The mean SSIM score between 2Ch view images and their reconstructions was 0.8053(+/- 0.0306), compared to 0.6330(+/-0.0389) for 4Ch view images. The 4Ch view encompasses larger atrial and ventricular regions, which likely contributes to its lower SSIM when 75% masks are applied. The mean PSNR score between 2Ch view images and their reconstructions was 22.3951(+/-1.9334), while that score was 16.9496(+/-2.1010) for 4Ch views images. The more complex structures in 4Ch view images result in a lower PSNR score. The whole evaluation metrics are shown in **Supplementary Table 4**.

The results demonstrated that the features extracted by the cineMAE successfully capture the majority of information present in the original MR images. We present several robust reconstruction examples alongside their corresponding originals in **Figure S2** and **Figure S3**. Notably, the reconstruction quality remains consistent and is not affected by changes in the ED and ES frame positions and both edge and texture information are well preserved.

### Interpretable model for mapping Heart-UDIPs to Heart regions

We first calculated the correlation between Heart-UDIP and 29 traditional heart related phenotypes based on multiple linear regression, we found that Heart-UDIP show significant correlation with them(P < 4.33e-6, F test, **Supplementary Table 6)**. In addition, as shown in Figure 1c, we developed our perturbation-based Decoder Interpretation (PerDI) framework to map Heart-UDIPs to the anatomical regions. We used 4Ch views for demonstration as they contain both left and right heart ventricles and atria. After extracting Heart-UDIPs from the input image, the original reconstructed images were generated from the decoder. To generate the perturbed images, we modified the Heart-UDIPs by adding one standard deviation to a certain dimension of interest. We performed the voxel-wise paired t-tests between the original and perturbed reconstructions across 500 individuals to identify the relevant regions for each dimension of interest.

The results were mapped to our heart atlas to show which anatomical regions each Heart-UDIP affects. Saliency mapping of the latent space reveals that individual feature dimensions demonstrate strong anatomical specificity, particularly in the end-diastolic frame. For instance, as shown in **Figure S4**, Heart-UDIP 28 primarily corresponded to the right ventricle, whereas Heart-UDIP 48 localized almost exclusively to the left ventricle. Heart-UDIP 248 corresponded to the right atrium while Heart-UDIP 19 covered the left ventricle and left atrium. This observation of overlapping but distinct features is consistent with the correlation analysis of the Heart-UDIP, which showed that the features were moderately correlated.

### Heritability of Heart-UDIPs

The Genome-wide Complex Trait Analysis (GCTA) software[13] was used to estimate the SNP-based heritability (h2) for each view of Heart-UDIP, defined as the proportion of variance explained by common autosomal variants. For the Heart-UDIPs at 2Ch view and 4Ch view, the SNP heritability was 0.14±0.07 and 0.15±0.06, respectively. Heart-UDIPs for 4Ch view showed slightly higher heritability than 2Ch view. On the other hand, stronger correlations were observed among the 2Ch Heart-UDIPs (absolute Pearson correlation coefficient |r| = 0.19±0.13) compared to the 4Ch Heart-UDIPs (|r| = 0.16±0.12). Additionally, the 4Ch view Heart-UDIPs included more comprehensive chamber information from CMR images, which might contribute to its higher heritability.

### Multivariate genome-wide association analyses of multi-view Heart-UDIPs

The Joint Analysis of GWAS (JAGWAS) software[9] was employed to perform two multivariate genome-wide association studies (mvGWAS) on 256 Heart-UDIPs derived from the 2Ch and 4Ch view (**Methods**). JAGWAS efficiently computes multivariate association statistics using single-phenotype summary data. The mvGWAS analyses encompassed 8,169,801 SNPs across the genome (**Methods**). Each SNP was evaluated for its association with Heart-UDIPs from multiple views, using a model that captures the distributed nature of genetic influences on the heart. To identify significant loci, we processed the mvGWAS results using FUMA[14], which performs LD-based clumping to define independent lead SNPs at each associated locus (r^2^ < 0.1). At genome-wide significance (P < 5×10^−8^), we identified 181 lead SNPs across 138 loci for the 2Ch view and 180 lead SNPs across 141 loci for the 4Ch view (**Figure 3, Figure S5**, and **Supplementary Table 7**). Comparison with the NHGRI-EBI GWAS Catalog[15] revealed substantial replication: 155 (85.64%) lead SNPs from the 2Ch view and 150 (83.33%) from the 4Ch view were previously reported (**Supplementary Table 8**). Notably, several identified SNPs are functionally relevant to cardiac biology (**Supplementary Table 9**). For example, rs11786896, an intronic PLEC variant, colocalizes with the atrial fibrillation GWAS signal and was prioritized as a candidate causal SNP[16], and rs6801957 locates in an enhancer region of SCN10A, influencing SCN5A expression—an essential gene for cardiac electrical conduction[17]. These results suggest that our proposed multi-view Heart-UDIP framework could effectively reveal the genetic underpinnings of heart function.

Additionally, 98 genomic loci were found to be shared between the 2Ch and 4Ch mvGWAS analyses, Notably, most loci identified in the 2Ch view (71.01%) were also detected in the 4Ch view. This likely reflects the broader anatomical coverage of the 4Ch view, which enables more comprehensive detection of heart-associated genetic loci. To further examine the shared phenotype architecture within different views, we performed canonical correlation analysis (CCA) to assess the variance explained between the 2Ch and 4Ch Heart-UDIP traits. The results aligned with the mvGWAS findings: the 4Ch Heart-UDIPs explained 23.35% of the variance in the 2Ch Heart-UDIPs, while the reverse direction accounted for 18.14% (Methods). This suggests the 4Ch view offers somewhat greater explanatory power for myocardium-related phenotypes. Interestingly, many overlaps lead SNPs from the two views demonstrated the important functional mechanism of heart development. For instance, rs17608766 (4Ch view, P = 3.317e-10, 2ch view, P = 5.06e-85) is located within the locus chr17:43,338,493– 45,936,354 near GOSR2—a gene previously associated with trabeculae carneae, critical structures in cardiac morphology[18] (**Figure S6**). Similarly, rs365990 (4Ch view, P = 1.30102e-95; 2Ch view, P = 8.736e-70) both located at chr14:23,861,811–23,899,027 near MYH6. —a gene encoding the α-heavy chain subunit of cardiac myosin, predominantly expressed in the atrial myocardium. Variants in MYH6 have been associated with atrial septal defects and atrial fibrillation, implicating its role in both cardiac development and electrical conduction[19-21]. (**Figure S6**). In addition, we found that several SNPs identified in the 4Ch view colocalize with known hypertrophic cardiomyopathy (HCM) associated loci (**Figure 2c & 2d**). For example, rs17617337, located in the BAG3 gene region, has been implicated in disrupting cardiomyocyte proteostasis and sarcomere stability[22]. Similarly, rs3176326 (6q21.2), an intronic variant in the CDKN1A locus, shows genome-wide significant association with sarcomere-negative HCM (P = 9.486e-45) and suggests a role for CDKN1A-mediated regulation of cardiomyocyte cell-cycle exit and hypertrophic growth independent of sarcomere mutations[5].

**Figure 2.**
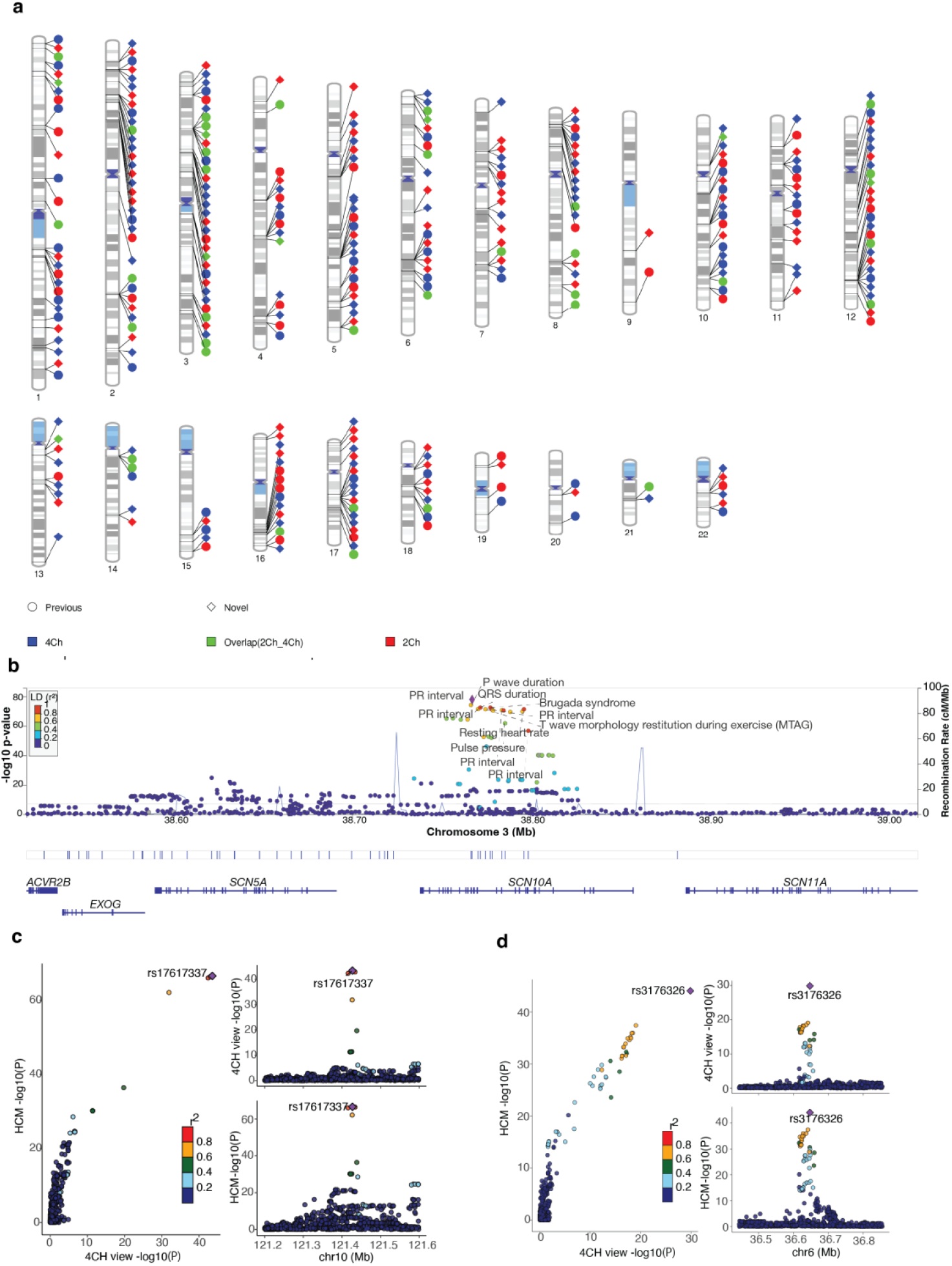
identified loci for 2Ch and 4Ch view mvGWAS. **(a)** Overlap between the identified loci and previous study. Colors indicate the view in which each SNP was identified blue for 4Ch, red for 2Ch, and green for SNPs common to both. Annotations highlight SNPs previously reported to be associated with heart function. **(b)** Common genomic region (3p22.2) between 2Ch and 4Ch view mvGWAS. **(c)** SNP colocalization between Heart-UDIPs and HCM at 10q26.11 region. **(d)** SNP colocalization between Heart-UDIPs and HCM at 6q21.2 region.

### Functional annotations of genetic loci associated with Heart-UDIPs

To investigate the biological function of significant SNPs identified for different view Heart-UDIPs, we used FUMA[14] to map those SNPs to genes through three strategies: physical position, expression quantitative trait loci (eQTL), and chromatin interactions (**Methods**). For the 2Ch view mvGWAS, we identified 524 unique genes. Of the 181 lead SNPs, 107 had at least one eQTL or chromatin interaction annotation, consistent with regulatory effects on gene expression by these variants or proxies in high linkage disequilibrium (**Supplementary Table 10, Figure S8**). For example, rs8082057 (P = 3.47e-08), located at chromosome 17p13.2, was identified as a regulatory variant for TRIB2, encoding a pseudokinase that modulates MAPK and AKT signaling, is expressed in cardiac tissues and has been implicated in cardiomyocyte stress responses and hypertrophic signaling pathways[23, 24].

For the 4Ch view mvGWAS, we identified 588 unique genes. Among 181 lead SNPs, 101 had at least one eQTL or chromatin interaction annotation (**Supplementary Table 10**). Notably, rs151041685 (P = 5.04×10-38), located at chromosome 2q31.2, was a significant chromatin interaction for TTN in left ventricle tissue. TTN encodes a giant sarcomeric protein essential for myocardial passive stiffness, sarcomere assembly, and cardiac contractility[25]. Furthermore, we test the gene set enrichment with some previously reported gene sets associated with the heart functions. We found both 2Ch and 4Ch views mvGWAS-identified gene set show significant enrichment for cardiac disease or related phenotypes (**Figure S7**).

### Gene-based association analysis and gene set enrichment analysis for Heart-UDIPs

Gene-based association analysis was performed using Multi-marker Analysis of GenoMic Annotation (MAGMA)[26], which integrates mvGWAS signals across SNPs mapped to each gene while accounting for linkage disequilibrium (LD). For the 2Ch view mvGWAS, we identified 318 genes reaching genome-wide significance (P < 0.05/19,291, **Supplementary Table 11, Figure S9**), 210 of which overlapped with genes previously annotated through at least one of three strategies: physical proximity, expression quantitative trait loci (eQTL) mapping, or chromatin interaction. These gene-based P-values were then used for gene-set enrichment analysis in MAGMA, referencing 15,488 curated functional gene sets from the MSigDB database[27]. 114 gene sets showed significant enrichment (Bonferroni-adjusted *P* < 0.05, Figure 3a, **Supplementary Table 12**). Most were related to cardiac development and structure, such asGO: heart development (*β* = 0.37, *P* = 2.65 e-18), GO: circulatory system development (*β* = 0.27, *P* = 2.10e-18, Wald test), GO: cardiac chamber development (*β* = 0.65, *P* = 8.76e-17), and GO: cardiac septum morphogenesis (*β* = 1.01, *P* = 4.892e-16, Wald test).

**Figure 3.**
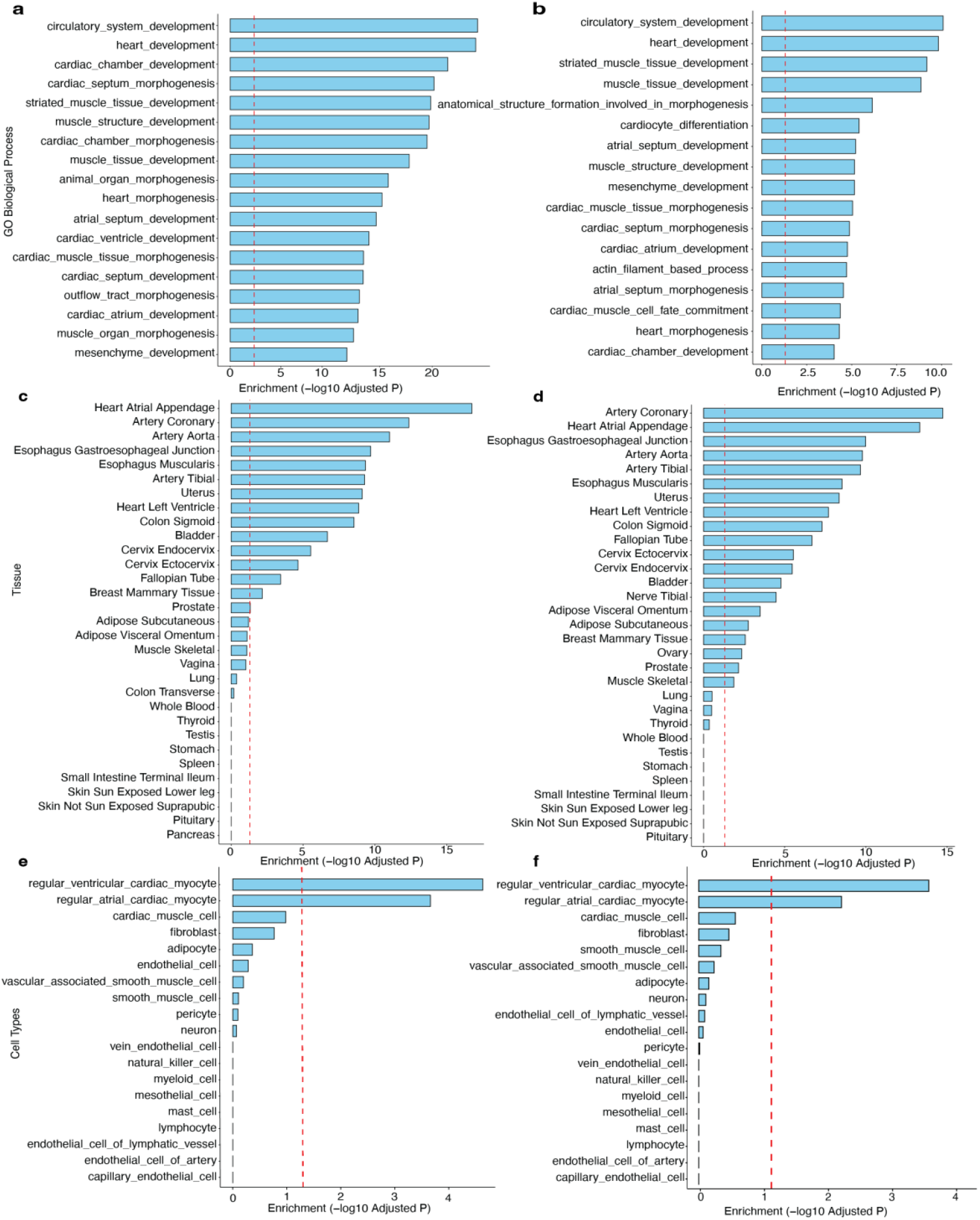
Gene-based association analysis for different view Heart-UDIPs. **(a)** Significant biological functional enrichment was observed for the 2Ch view of the Heart-UDIPs. **(b)** Significant biological functional enrichment was observed for the 4Ch view of the Heart-UDIPs. **(c, d)** Tissue-specific enrichment of genes identified from the 2Ch and 4Ch views of the Heart-UDIPs. **(e, f)** Cell type–specific enrichment of genes identified from the 2Ch and 4Ch views of the Heart-UDIPs. The red dashed line indicates the Bonferroni-adjusted significance threshold (P < 0.05).

For the 4Ch view mvGWAS, we identified 288 genome-wide significant genes (*P* < 0.05/19,291; **Supplementary Table 11, Figure S9**), of these, 218 overlapped with genes annotated by at least one of three mapping strategies from FUMA. MAGMA gene-set analysis identified 80 significantly enriched sets (Bonferroni-adjusted P < 0.05 across 15,488 sets, **Supplementary Table 12; Figure 3b**), largely overlapping those from the 2Ch-view analysis and highlighting functions related to cardiovascular development. Notably enriched terms included GO: cardiocyte differentiation (*β* = 0.52, *P* = 2.30e-10, Wald test; the most significant) and GO: atrial septum development (*β* = 1.291, *P* = 3.59e-10, Wald test).

To investigate tissue- and cell-type-specific regulatory relevance, we first examined the expression specificity of associated genes across 54 tissues from GTEx v8[28]. Both Heart-UDIPs associated genes for 2Ch and 4Ch view showed significant upregulation in six cardiac-related tissues after Bonferroni correction (**Figure 3c and 3d**). For example, in the heart atrial appendage (2Ch: *β* = 0.11, *P* = 3.37e-19; 4Ch: *β* = 0.09, *P* = 9.02e-16), artery coronary (2Ch: *β* = 0.11, *P* = 8.14e-15; 4Ch: *β* = 0.11, *P* = 3.38e-17), Artery Aorta (2Ch: *β* = 0.06, *P* = 3.23 × 10^−8^; 4Ch: *β* = 0.05, *P* = 6.69 × 10^−7^), and Heart Left Ventricle (2Ch: *β* = 0.07, *P* = 2.50e-11; 4Ch: *β* = 0.06, *P* = 3.75e-10), expression enrichment was statistically significant. We further evaluated gene-based associations in a single-cell gene expression dataset from twenty-five human heart samples, including six heart tissues[29]. Both Heart-UDIPs associated gene for 4Ch and 2Ch view showed significant upregulation (*P* < 0.05/19) in heart-related cell types(**Figure 3e and 3f**), including regular atrial cardiac myocytes (2Ch: *β* = 0.07, *P* = 1.95e-6; 4Ch: *β* = 0.09, *P* = 6.95e-7; Wald test) and regular ventricular cardiac myocytes (2Ch: *β* = 0.09, *P* = 2.10e-7; 4Ch: *β* = 0.07, *P* = 2.51e-5; Wald test).

### Heart diseases relevance of Heart-UDIPs

We evaluated the discriminative performance of Heart-UDIPs by classifying six cardiovascular diseases (**Supplementary Table 5**). These patients are picked up by the ICD-10 codes. For each disease, we randomly choose the same number of cases from the control group. We calculate the evaluation matrix of AUC for each of them. The ablation study results from this table show the efficiency of our masking strategy. By forcing the cineMAE to reconstruct the images with missing values, the model extracts more compact UDIPs from the input images. For most of these diseases, Heart-UDIP with masks outperforms the regular ones in both 2Ch and 4Ch views.

The results show that our extracted features have predictive power across a range of cardiovascular and metabolic disorders. The AUC scores (0.7992 (+/- 0.0215) and 0.8084 (+/- 0.0233) of Heart-UDIP with mask) in heart failure and atrioventricular block prediction indicate that features effectively capture structural and functional biomarkers relevant to impaired cardiac conduction and pump function. The model also achieved a high AUC score of 0.8921 (+/- 0.0626) in HCM and 0.9444 (+/- 0.0333) in DCM with the 2Ch Heart-UDIP with mask. These results suggest the sensitivity of our feature set in detecting nuanced morphological patterns associated with HCM, a disease often challenging to diagnose in early stages.

Beyond the robust classification results, our method shows potential for uncovering latent signatures across a spectrum of myocardial diseases. The consistent performance on atrial fibrillation[30] and myocardial infarction[31] suggests our features encode both structural abnormalities and potential rhythm-related alterations. The features’ ability to differentiate diabetic patients suggests a link between cardiac imaging biomarkers and systemic metabolic health—opening the door for early detection of comorbidities through routine cardiac scans. These findings support the use of our feature set not only for disease classification but also for hypothesis generation in cardiovascular research and personalized risk stratification.

### Heart-UDIPs show significant association with heart related diseases

Polygenic risk scores (PRS) provide an estimate of an individual’s genetic predisposition to specific traits. Using PRS-CS[32], we computed PRS for 57,867 individuals in the UK Biobank cohort across seven heart-related traits derived from FINNGEN [33] (**Methods**): cardiac rhythm, complex congenital heart defects, ischemic heart disease, major coronary artery disease (CAD), and hypertension, hypertrophic cardiomyopathy (HCM), dilated cardiomyopathy(DCM). We then assessed the relationship between multi-view 256-dimensional Heart-UDIPs and these PRSs using a multivariate association model (Methods; **Figure 4a**). In both 2Ch and 4Ch views, Heart-UDIPs showed significant associations with cardiac disease, such as HCM (P < 7.96e-162, F-test), hypertension (P < 3.43e-88, F-test), DCM (P < 7.96e-162), indicating that the Heart-UDIPs capture genetic risk linked to clinically relevant cardiac conditions.

**Figure 4.**
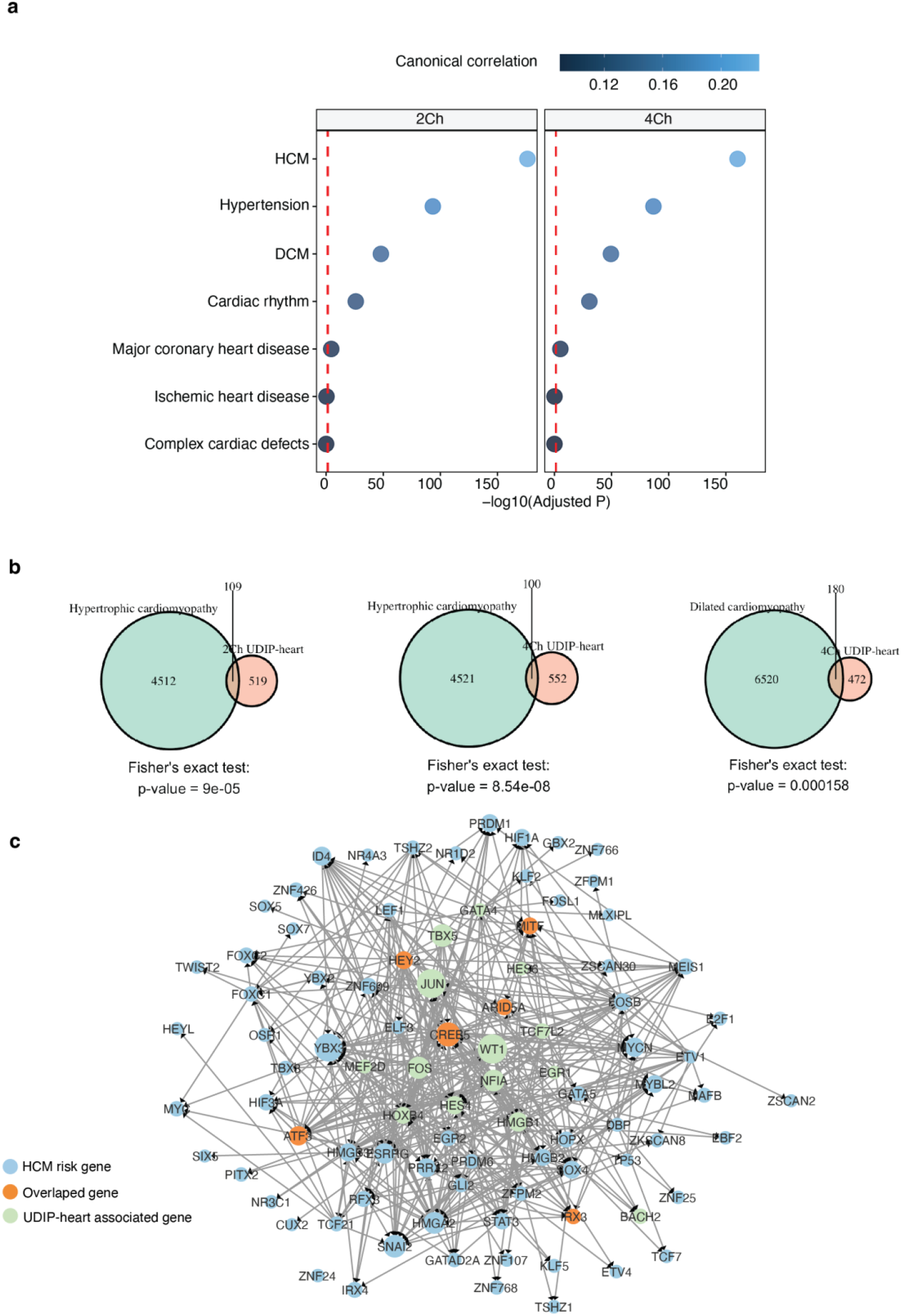
Association between hearts related disease and Heart-UDIP from different views. **(a)** Association between Heart-UDIPs and the PRS score for different cardiac diseases. **(b)** Significant gene set enrichment between Heart-UDIPs related genes and risk genes for HCM and DCM. **(c)** Biological regulation between Heart-UDIPs related gene and HCM risk gene in the heart specific transcriptional regulatory network. The edge in this network means the regulation from transcription factor to the target gene.

On the other hand, we explored the enrichment of identified Heart-UDIPs related genes in different views with 39 cardiac disease-related risk gene set. After Bonferroni correction (0.05/39), we found that both 2Ch and 4Ch were significantly enriched for many cardiac diseases (**Figure S10**), especially in HCM (P < 9e-5, Fisher’s exact test, **Figure 4b**) and atrial fibrillation (P < 5.66e-15, Fisher’s exact test, **Figure S10**). Additionally, 2Ch was enriched for aortic valve insufficiency (P = 6.49e-4, Fisher’s exact test), which was primarily associated with the left ventricle, which develops progressive dilation and functional decline secondarily. The 4Ch was enriched for DCM (P = 1.58e-4, Fisher’s exact test). Interestingly, some genes identified by both the 2Ch and 4Ch view are risk genes for a variety of cardiac diseases. For example, TCF21, a basic helix-loop-helix transcription factor, governs the differentiation of cardiac fibroblasts during development and regulates fibroblast activation in adult hearts, and genetic and functional studies have shown that TCF21 modulates coronary artery disease and atherosclerosis risk by promoting vascular smooth muscle cell phenotypic switching and stabilizing atherosclerotic plaques[34-36].

Having previously found a significant association between Heart-UDIP and HCM, we further explored the regulatory relationship between Heart-UDIP related genes and HCM risk genes in a heart-specific regulatory network(**Figure 4c**) [37]. Notably, Heart-UDIP related transcription factors such as *WT1, TBX5*, and *FOS* form distinct regulatory axes that converge on key HCM risk genes. For example, *WT1* functions as a transcription factor reactivated during cardiac injury and remodeling[38], its regulatory influence extends to CREB5, and in a mouse model of pressure overload-induced heart failure, *CREB5* has been shown to mediate cardiomyocyte energy metabolism and attenuate myocardial remodeling through the Creb5/NR4a1 axis in the PI3K/AKT pathway, downstream of GLP-1 receptor signaling [39]. This indicates a *WT1* → *CREB5* axis, suggesting that WT1-driven CREB5 activation may contribute to pathological hypertrophic pathways and fibrotic remodeling observed in HCM.

### Comparison with GWAS on other MR heart image derived phenotypes

To compare our findings, we reviewed previous GWAS studies of heart imaging-derived phenotypes based on UKBB CMR data in **Table 1**. These included three studies using high-latitude CMR-derived phenotypes (Sooknah et al.,[40]; Burns et al.[41]; Bonazzola et al.[42]) and three using traditional CMR-derived phenotypes (Meyer et al.[18]; Pirruccello et al.[3]; Aung et al.[1]) from short axis(SA) view images. We compared our results across multiple levels. Our study is the first to jointly assess the genetic discovery potential of both LA 2Ch and 4Ch Heart-UDIP views. This different view framework allowed us to identify a broader spectrum of associated SNPs and genes, with several findings replicated across existing studies (**Figure S11**). For example, *TTN*, which encodes the largest sarcomeric protein vital for cardiac contractility and structural integrity, was consistently detected in all five prior studies. We also replicated findings related to *CDKN1A* (p21), a regulator of cardiomyocyte senescence and remodeling though the p53 pathway under cardiac stress [43]; and *GOSR2*, a vesicle trafficking gene recently linked to mitral valve morphology and cardiac structure [44], both of which appeared in three other studies.

**Table 1.**
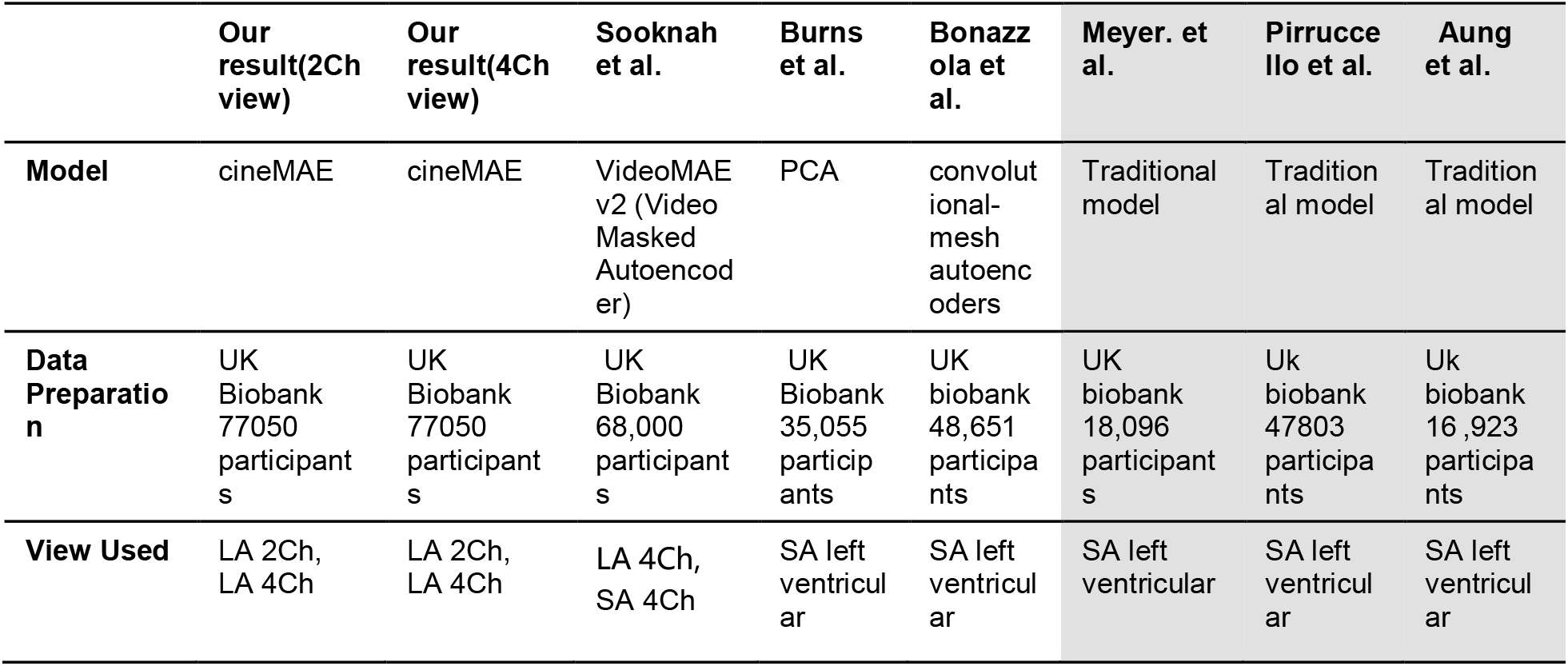
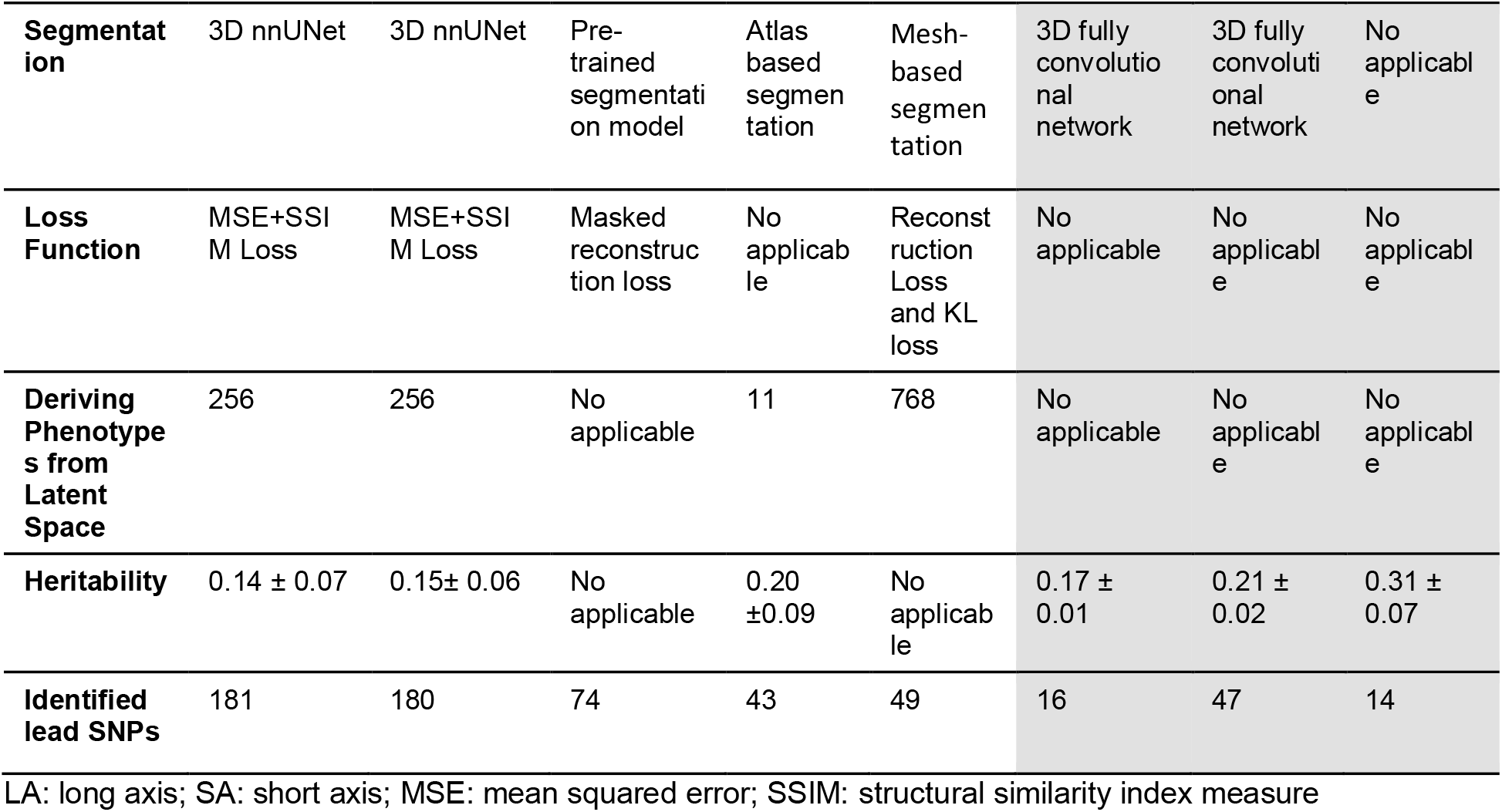
Comparisons between our Heart UDIP-based genetic discovery and other published ones.

Additional genes such as STRN, involved in PP2A signaling, cardiac arrhythmia, and remodeling [45], *BAG3* (a sarcomere integrity regulator via chaperone-assisted autophagy, associated with DCM)[46], and *SMG6* (a key player in nonsense-mediated mRNA decay with possible roles in cardiac gene regulation)[47], were detected in two additional studies.

Moreover, our analysis uncovered some novel genes. For instance, *ADAMTS18*, a secreted metalloproteinase involved in extracellular matrix remodeling, has not been widely studied in cardiac function. However, in mice, ADAMTS18 is expressed around the aortic arch and carotid arteries during embryogenesis. Its deficiency leads to vascular malformations and impaired vascular smooth muscle differentiation via Notch3 pathway disruption [48].

Another novel gene, *ARNT*, a transcription factor that partners with HIF-α subunits, plays a crucial role in cardiac metabolism. Cardiac-specific ARNT deletion in mice causes lipid accumulation and defective fatty acid oxidation, resulting in cardiomyopathy and highlighting its role in metabolic homeostasis [49].

### Summary of our cineMAE model

We present a novel framework using a 3D U-architecture autoencoder (cineMAE) to learn deep image phenotypes from CMRIs for genetic discovery. The whole system consists of data preprocessing, model training, feature interpretation and GWAS analysis. Several innovative contributions have been made to find new heart related significant SNPs. Specifically, the data preprocessing pipeline helps to exclude those irrelevant organs, resulting in computational efficiency and feature concentration. The cineMAE is able to reconstruct high-quality images with a mask ratio of 75%. This demonstrates that there is much redundant information in those CMRIs. By removing those redundancies with our masking strategy, our Heart-UDIP, with much fewer feature correlation score, is able to find a greater number of significant SNPs in the GWAS analysis.

We validated our Heart-UDIP through a perturbation-based interpretation method. The saliency map of the specific Heart-UDIP in our heart atlases shows the relationship between the Heart-UDIPs and the anatomical structures. Our interpretation method show that individual features are mapped to specific anatomical regions of the heart, including the ventricles and atria.

Our unsupervised cineMAE is good at learning heart function related information which can be proved in both image feature evaluations and genetic discoveries. For example, the seven heart disease classification results show that heart disease-related information is well captured in our Heart-UDIP. Meanwhile, some of the found SNPs are also linked to those diseases. The consistency in those discoveries proves that our Cine MAE is efficient in learning critical information from CMRIs.

This strong concordance between our imaging-based interpretation method and genetic discovery demonstrates that the unsupervised cineMAE framework is highly effective at extracting clinically and biologically meaningful information. This approach holds significant potential for accelerating imaging genetics research across other organs.

## Discussion

In this study, we extended our previously developed UDIP framework and developed cineMAE for cine cardiac MRI feature extraction, enabling genetic association studies without relying on predefined clinical phenotypes or manual image annotations. By applying this framework to cardiac magnetic resonance images from the UK Biobank, we successfully derived robust, heritable, and interpretable imaging phenotypes (Heart-UDIPs) from unsupervised deep learning models trained on 2Ch and 4Ch view CMRIs. Our multivariate GWAS on these Heart-UDIPs identified numerous genetic loci, including novel SNPs associated with cardiac function and structure, significantly expanding the known genetic architecture of the heart. Moreover, our integrative analysis demonstrates that the Heart-UDIPs shows significant association with various cardiac diseases and reveals the potential key regulatory pathways from Heart-UDIP-related genes to the risk gene of HCM. These findings highlight the power of unsupervised deep learning methods for uncovering genetic and molecular insights into cardiac diseases.

Our cineMAE model has successfully tackled the challenges brought by cine MR images. The first challenge comes from the redundant 2D structure information across the long-axis images. Meanwhile, the pattern of dynamic changes in the ventricle area is an important indicator of heart functions. Thus, a compact representation of the cine CMR with discriminative information is needed for subsequent analysis. The second challenge is from the readiness of the cine CMR. Different from the brain MRI in the UKBB that were extensively processed via a well-established pipeline, there are no segmentation masks available for CMR. The surrounding organs in the CMR introduce irrelevant information to the imaging phenotypes. Additionally, there are no registration atlas for CMR of UKBB which is a critical component for GWAS analysis. To significantly reduce the preprocessing burden, we developed a uniform preprocessing pipeline for Cine CMR images, summarized as follows.

The purpose of heart segmentation is to eliminate the signal from other organs in heart related SNP discovery with our Heart-UDIP. We only annotated 100 cases for both 2Ch-view and 4Ch-view to finetune the nnUNet model as it has robust segmentation across different segmentation tasks. After fine-tuning, we use it to process all the 2Ch and 4Ch CMR images in our dataset with more than 75000 samples. With the segmentation results, we crop the original input images into the resolution of 80*80*50 for the following processing.

The primary purpose of heart image registration is to align the CMR images of different individuals to a common space. There is no atlas available for heart image registrations, which is a critical step in feature extraction. Thus, we built the heart atlas based on 150 randomly selected cases with the ANTs Python package. The models are trained on the registered images with resolution of 80*80*50.

As a result, our approach significantly reduces the burden on preprocessing compared to conventional methods. It does not require manual or automated segmentation of regions of interest within the heart, such as the ventricles and great vessels which are widely used in traditional image feature-based genetic discoveries. Furthermore, it bypasses the complex step of 3D mesh construction, simplifying the workflow to a single image format transformation.

To enhance the power of our genetic discovery pipeline, we introduced a masked reconstruction strategy during model training. This approach was motivated by our observation that our Heart-UDIPs, which are from the models trained with full data, exhibited high inter-feature correlation, significantly higher than those from our brain UDIP. This redundancy can diminish the statistical power of subsequent GWAS analyses. Therefore, we trained our network with randomly masked-out images. The resulting Heart-UDIPs show a significant reduction in correlation scores, indicating that the masking strategy successfully forced the network to learn more discriminative and information-rich features, providing a more robust basis for identifying novel genetic loci.

**Figure S**2 and **Figure S3** show that these models capture the whole series information of the heart images, as the reconstructed images show similar content to the original one. The dynamic ventricle area changes between the original images and the reconstructed images are consistent, which means that our image features capture both the texture and spatial information of the original input.

Exploring the genetic architecture of the heart using Heart-UDIPs, we identified 163 genetic loci across multiple anatomical views, including 75 novel loci previously unassociated with cardiac traits. Notably, rs1763607, located near HSPB7—a small heat shock protein abundantly expressed in cardiac tissue— has been linked to cardiomyopathy. Functional studies in mice have demonstrated its role in maintaining sarcomere integrity and supporting the cardiac muscle’s response to stress. Likewise, rs35001652 is situated near KDM1A, a lysine-specific histone demethylase. Current evidence indicates that KDM1A plays a regulatory role in heart development, with gene disruption leading to structural heart defects in animal models[42]. Additionally, rs11264422 lies proximal to CHRM2, which encodes the M2 muscarinic acetylcholine receptor. This receptor modulates parasympathetic tone and resting heart rate, with genetic variants linked to altered heart rate variability and increased arrhythmia risk in humans[50].

Our study further elucidates the relationship between Heart-UDIPs and HCM, demonstrating how unsupervised deep learning-derived phenotypes capture nuanced yet biologically meaningful variations in cardiac morphology and function relevant to HCM pathogenesis. Heart-UDIP phenotypic features closely resemble clinical manifestations of HCM, suggesting their potential utility for disease classification. Furthermore, we identify a robust predictive link between genetic variants associated with Heart-UDIP and HCM risk. At the molecular level, colocalization analyses reveal significant overlap between SNPs identified through Heart-UDIP genome-wide association studies (GWAS) and previously established HCM genetic loci, reinforcing the genetic and biological significance of these findings. Additionally, the JUN gene, encoding the AP-1 transcription factor c-Jun, occupies a central position within the regulatory network governing HCM. JUN directly interacts with key transcriptional regulators, including FOS, CREB, and WT1, highlighting its role as a critical nodal coordinator of hypertrophic pathways. Mechanistically, JUN integrates pathological stimuli through JNK/MAPK signaling pathways, forming c-Jun, Fos heterodimers that directly influence gene expression associated with cardiac hypertrophy, fetal cardiac gene activation, fibrosis, and inflammation, partially via cross-activation with CREB [51, 52]. Functional experiments further illustrate that cardiomyocyte-specific deletion of Jun disrupts sarcomere protein expression, exacerbates interstitial fibrosis, and promotes maladaptive cardiac remodeling under conditions of pressure overload[53]. Collectively, these findings position c-Jun as a master transcriptional regulator, orchestrating structural and stress-responsive gene programs central to HCM pathogenesis.

As shown in **Table 1**, we compared different approaches on heart imaging-based genetic discoveries. Three common deep learning architectures which are 3DCNN, 3DViT and 3D mesh networks are included in this table. Our cineMAE has a 3D U-shape architecture with three encoder blocks and decoder blocks. Compared with 3D ViT models, ours requires fewer training samples and computational resources. With our architecture, the MAE-based masking strategy helps the Heart-UDIP to find more significant SNPs (**Supplementary Table 1**). Those Heart-UDIP also demonstrate their quality in disease classification tasks (**Supplementary Table 2**). Unlike the 3D mesh networks, we don’t have to convert the original MRIs to other formats with fundamental difficulties in implementing, which simplifies the data preprocessing step in our task. In **Supplementary Table 1**, we can see that our data preprocessing pipeline also contributes to the genetic discoveries. Although the mean heritability of Heart-UDIP was lower than that reported in some previous studies, these phenotypes comprise many distinct dimensions with relatively low inter correlation. This suggests that genetic variance may be distributed across dimensions, with each dimension capturing partially independent components of the overall heritability and potentially reflecting distinct genetic influences. To more fully harness the multidimensional nature of Heart-UDIP, we applied a multivariate GWAS framework that jointly integrates information across dimensions. This approach increases statistical power, yielding a greater number of genome-wide significant loci and enhancing the discovery of genetic factors underlying cardiac structure and function.

## Methods

### Data preprocessing pipeline

Our data preprocessing pipeline comprises six sequential modules designed to convert raw DICOM images into standardized, analysis-ready volumes. The initial step converts all DICOM series into the NIfTI format. Subsequently, every NIfTI file undergoes spatial registration to align it within our heart atlas, followed by cropping to a uniform resolution of 80 x 80 x 50 voxels. It is important to note that the corresponding image annotations are processed separately and used exclusively for training the nnUNet segmentation model. Further details on the specific registration and segmentation methods are provided in the following sections.

### Image segmentation with nnUNet

Whole-slice images with a resolution of more than 208*168*50 contain surrounding anatomical structures and background noise, which can introduce irrelevant features and hinder the network’s ability to focus on learning meaningful representations of the heart. To reconstruct the whole slice images, more parameters are needed, which increases the burden of 3D network training. Additionally, the features extracted from those slices may lead to the discovery of loci unrelated to heart structure and function. Thus, heart segmentation is a crucial step in our data pipeline for both building the heart atlas and training our 3D image reconstruction networks.

In our work, we annotate 100 cases for both 2Ch and 4Ch view images. They are utilized to train the nnUNet[12] corresponding to their view. After training converges, the nnUNet models are deployed to segment our whole heart dataset. Based on the segmentation results, we crop the image to a resolution of 80*80*50 for those two view images. The cases with no heart segmentation masks by the nnUNet are excluded from the whole heart dataset.

### Heart atlas building with ANTs

As our cineMAE is trained in an unsupervised way, image registration is a critical constrain on the training data to ensure spatial correspondence across all those 77,050 patients. It will help the Cine MAE to learn high-level, spatially-aware features that represent meaningful anatomical phenotypes.

We build our Heart atlas with ANTs for both 2Ch view and 4Ch view images. For each view atlas, we randomly pick up 150 Cine MRIs as the training dataset. Based on the segmentation results of those Cine MRIs, we crop them to a resolution of 80*80*50 where the heart areas are well captured. We use ANTsPy package to create the atlas from those 150 cropped Cine MRIs. To avoid blank slices caused by z-axis shift, we add the transformation restriction during the registration process in the template building. The template building process has three iterations of registering training images to the final template. After that, we save the template as the atlas for Cine MRIs registration.

### Image registration with ANTs

Image registration[54] is an important tool to find correspondence among different images. With that information, deformations in medical images both spatially and temporally can be modified. As a result, deep learning models can find better high-level image features for subsequent tasks, like image segmentation and classification.

Since there is no whole-heart atlas available for heart MRI registration, we constructed a template using images from 150 patients with ANTs[55]. To assess the quality of this template, we employed similarity metrics such as Structural Similarity Index (SSIM)[56], Mutual Information (MI)[57], and Normalized Cross-Correlation (NCC)[58] to quantify the similarity between the template and the original images. As a control, we also measured these metrics using a randomly selected case as the reference to compare its performance against the template.

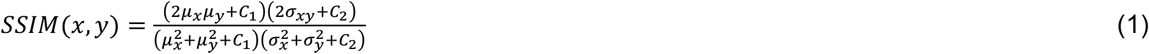

where µ is the mean intensity and σ is the standard deviation, and C is a constant value.

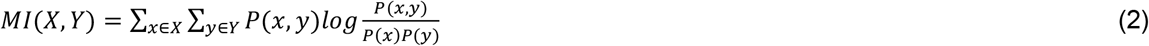

where P is a probability distribution.

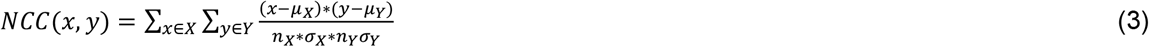

Where *n*_*X*_ is the number of image pixels in image X and *n*_*Y*_ is the number of image pixels in image Y. µ and σ are the average and standard deviations of image.

When using the template as a reference, the average SSIM between the template and other cases is 0.483, the MI is 0.612, and the NCC is 0.709. In contrast, when a randomly selected case is used as the reference, the SSIM drops to 0.259, MI to 0.211, and NCC to 0.317. These significant improvements indicate that the template provides a more structurally consistent and representative reference for aligning different heart images, leading to better similarity measures across cases.

### Implementation of our image reconstruction model

As shown in **Figure 1b**, we employ a 3D convolutional masked autoencoder framework to learn the UDIP of cardiac images. The network is designed as a lean autoencoder with three encoder layers and three decoder layers connected by a bottleneck layer. It is trained on MSE loss and weighted SSIM loss between the original images and their corresponding reconstructions. The loss functions are listed below.

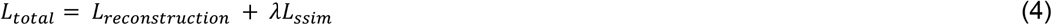

Each encoder block comprises a 3D convolutional layer, an instance normalization[59] layer and ReLU[60] layer. Given the relatively small batch size used during training, instance normalization is chosen to enhance the quality of the generated images. A skip connection is introduced between the last encoder block and the first decoder block to mitigate blurring in the reconstructed images. To balance information flow, the weighted parameters for these blocks are set to 0.2 and 0.8, ensuring that most of the reconstruction-relevant information comes from the bottleneck layers.

As there are many redundant 2D structure information in the Cine MRIs. In order to get more compact image features, we apply the masks to all those 50 frames so that the cineMAE has to learn to reconstruct the missing structures with the rest of the images. The masks, with a default ratio of 75%, are generated randomly to cover different positions of the images.

The 3D network is trained for 200 epochs with an initial learning rate of 1e-4. Model parameters are optimized using the Adam optimizer, with a batch size of 4, on an NVIDIA A100 GPU. The dataset is split into training and testing subsets in a 4:1 ratio.

We evaluate the quality of the generated images from our generator by the structure similarity (SSIM)[56] and peak-signal-to-noise ratio (PSNR)[61].

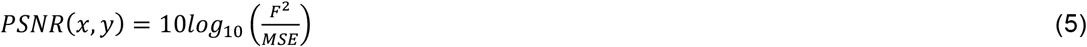

Where F is the function to get the maximum fluctuation of the pixel values of the image.

### Perturbation-based Decoder Interpretation for Heart-UDIPs

In this manuscript, we used the perturbation-based decoder interpretation (PerDI) framework to show the saliency map of our Heart-UDIPs on the 2D+T cine MRIs. For each dimension of Heart-UDIPs, we added one standard deviation to their original values and used the decoder to get a perturbed reconstruction. The difference between the original reconstruction and the perturbed reconstruction was then used as an indicator of the saliency map of the perturbed Heart-UDIP. T-statistics between the original reconstructions and their corresponding perturbed reconstructions among 500 randomly selected patients were calculated. Those results were mapped to our heart atlas where the areas with higher absolute values are considered as the area most influenced by a specific Heart-UDIP. The first frame of several 4Ch view patients’ perturbation-based results are shown in **Figure S4**.

### Heart-UDIPs-based Disease Classification

We select the patients with their ICD-10 codes and pick up the same number of patients as the control group. Their corresponding Heart-UDIPs from our cineMAE are utilized to train a logistic regression model. Five-fold evaluations with AUC score are utilized to test the heart related disease classification with our Heart-UDIPs. For each disease, we compute the five-fold AUC scores for 2Ch view Heart-UDIPs, 4Ch view ones and their combinations, as shown in **Supplementary Table 5**. It helps us to determine whether different view Heart-UDIPs capture unique information in their view. The results are also utilized to measure the enhancement of Heart-UDIPs caused by our masking strategy (**Supplementary Table 2**).

### Genotype data preprocessing

Genetic analyses were performed on individuals from the UKB with European ancestry, who also had genotypes and CMR images data. Standard quality control procedures were then applied to the UKB v3 imputed genetic data[62]. These procedures included the following steps: (1) exclusion of individuals with failed genotyping, abnormal heterozygosity status, or withdrawn consents. (2) removal of participants genetically related—up to the third degree—to another participant, as inferred by kinship coefficients implemented in PLINK[63]; (3) elimination of variants with a minor allele frequency below 1%; (4) removal of variants with a genotype missing rate exceeding 10%; (5) exclusion of individuals who were part of the deep learning model’s training process to prevent data leakage and ensure independent validation. After quality control, we retained 57,867 individuals and 8,925,988 variants. The Hardy-Weinberg equilibrium test was not conducted: although it is a standard practice for QC genotype data from directly genotyped markers, it may not be statistically sound to convert the dosage back to the best guess imputed genotypes and do HWE again.

### GWAS analysis for Heart-UDIPs

For the GWAS analysis, we first used fastGWA from the GCTA package (Version 1.94.1) to run linear mixed model association tests on the 512 embeddings obtained from the 2Ch (256 dimensions) and 4Ch (256 dimensions) CMRIs, adjusting for several covariates. These covariates included age (field ID: 21003), age squared (a^2^), sex (field ID: 31), sex × age, sex × age^2^, BMI(field ID: 21001), genetic ethnic grouping (field ID: 21006), Ethnicity (field ID: 21000), the first 10 genetic principal components (field ID 22009), scanner table position (field ID 25759), assessment center location (field ID 54), and date of assessment (field ID 53).

We also accounted for kinship information provided by the UK Biobank. Following the single-variant GWAS, we used JAGWAS[9] to combine the results into multi-variant association analyses across the 256 UDIP-heart from the 2Ch and 4Ch views. A genome-wide significance threshold of P < 5×10^−8^ was applied for SNP-UDIP associations in both views.

### Identification of genomic loci, functional annotation, and gene annotation of Heart-UDIPs

We used FUMA (v1.70) [14] to identify genomic loci with significant multivariate associations with UDIP-heart and to perform functional annotation using default settings. Linkage disequilibrium (LD) was defined using the European panel from the 1000 Genomes Project. SNPs reaching genome-wide significance (mvGWAS P < 5 × 10^−8^) and showing moderate LD (r^2^ < 0.6) with other significant variants were first identified. Additional SNPs in high LD (r^2^ ≥ 0.6) with these were then included for annotation.

Independent lead SNPs were defined as those with low LD (r^2^ < 0.1) relative to other significant SNPs. Genomic loci were constructed by merging LD blocks of significant SNPs located within 250 kb of each other, allowing for multiple independent lead SNPs within a single locus. The MHC region was excluded due to its complex LD structure. Functional annotation and SNP-to-gene mapping followed FUMA protocols as described in prior studies[41].

### Gene set enrichment analysis

Gene-set enrichment was assessed using MAGMA (v1.08), as implemented in FUMA with default settings. This analysis tested whether gene-based P-values for 19,291 genes were lower within predefined functional sets compared to the rest of the genome, accounting for confounding factors such as gene size and SNP count. A total of 15,488 gene sets from MSigDB v7.0[64] were analyzed, including 5,500 curated sets, 7,343 GO biological processes, 1,644 GO molecular functions, and 1,001 GO cellular components. Bonferroni correction was applied to control for multiple testing (P < 0.05/15,488), separately for each mvGWAS. The most significant GO biological processes were selected for visualization in the figure.

### Gene expression analysis

To evaluate the tissue and cell-type specificity of UDIP-heart related genes, we analyzed expression data from 54 tissues in the GTEx dataset. Using a linear regression model, we applied MAGMA to test for significant enrichment of genes highly expressed in specific tissues. For cell-type analysis, we used the CELL TYPE function in FUMA, which tested whether gene-based association z-scores were positively correlated with expression levels in single-cell RNA sequencing data from the adult human heart. This data included six heart tissues and 19 cell types. Associations were considered significant if the P values passed the Bonferroni-corrected threshold for the number of tissues (P<0.05/54) or cell types(P<0.05/19) tested.

### Polygenic Risk Score of heart associated diseases

The polygenic score (PGS) represents an estimate of an individual’s genetic risk for a given trait. To estimate the association between Heart-UDIPs at different views and genetic risk for different heart-related traits, the polygenic score for six complex traits was generated using PRScs with default parameters. To ensure the robustness of the PRS results, we selected GWAS studies that did not overlap with the UKB sample that we used in this study. We used six complex traits obtained from three separate sources. We analyzed seven complex cardiovascular traits drawn from three independent sources. Five traits—cardiac rhythm, complex congenital heart defects, ischemic heart disease, major coronary heart disease (CHD), and hypertension—were obtained from the FinnGen GWAS. Dilated cardiomyopathy (DCM) was from Zhang et al. [4] and hypertrophic cardiomyopathy (HCM) from Tadros et al.[5].

To examine the relationship between multiple Heart-UDIPs and disease-specific PRSs, we used canonical correlation analysis (CCA). First, we applied linear regression to remove the effects of covariates (adopted for GWAS analysis) from the Heart-UDIPs. We then calculated correlations between the residuals of the Heart-UDIPs and the PRSs for various diseases. Finally, we applied Bonferroni correction and retained only the associations with statistically significant P values (P < 0.05).

## Author contributions

DZ and JN initialized the conceptualization of the project. LY, ZX, and DZ designed the overall methodology. LY leads the development of data preprocessing pipeline and cineMAE for this paper. XZ leads the post GWAS analysis for the Heart-UDIP. C.Chen fine-tunes the nnUNet for Cine MRIs segmentation. KM and ZX are in charge of running GWAS analysis for the Heart-UDIP. DZ, ZX, LY, and XZ interpreted the results. LY and XZ developed the majority of the initial draft of the manuscript. C.Cassidy, KP and DZ contributed to the writing of the manuscript. All authors reviewed and approved the manuscript. EA, JN and DZ are the project leaders and funders.

## Data availability

The data used in this study is derived from the UK Biobank under application 24247. Researchers interested in replicating or extending our work can apply for access through the official UK Biobank website.

The complete GWAS summary statistics have been deposited in the GWAS Catalog (accession GCP001376).

## Code availability

Upon publication of our paper, we will release all source code used in our experiments, including data preprocessing, model training, evaluation scripts and the model weights, in both GitHub and Hugging Face repository.

## Acknowledgements

This research is funded by U01AG070112, R01EY032768, K. Lance Gould Distinguished University Chair in Coronary Pathophysiology.

